# Metabolic exchange and siderophore sharing underlie emergent biofilm synergism

**DOI:** 10.64898/2026.02.24.707706

**Authors:** Jiyu Xie, Xinli Sun, Viktor Hesselberg-Thomsen, Kun Duan, Huihui Zhang, Weibin Xun, Nan Zhang, Mikael Lenz Strube, Qirong Shen, Ákos T. Kovács, Ruifu Zhang, Zhihui Xu

**Author notes:** Corresponding authors: Zhihui Xu, Ákos T. Kovács, and Ruifu Zhang, (Z.X.), (A.T.K.), (R.Z). Jiyu Xie and Xinli Sun contributed equally to this work.

## Abstract

Biofilm communities exhibit emergent properties that exceed the sum of contributions from individual members of the community. Here, we describe a multilayered metabolic interaction that drives enhanced biofilm formation among three bacterial species from the plant rhizosphere. Comparative metatranscriptomic and metabolomic analyses reveal that *Bacillus velezensis*-secreted 5-aminovaleric acid promotes the growth of the other community members, *Burkholderia contaminans* and *Acinetobacter baumannii*. In return, *B. contaminans* supplies branched-chain amino acids for *B. velezensis*. Branched-chain amino acids and cell–cell signaling acyl-homoserine lactones from *B. contaminans* induce biosynthesis of the siderophore bacillibactin in *B. velezensis*, that is further enhanced by *A. baumannii*. In exchange, the *B. velezensis*-secreted siderophore promotes the growth of *B. contaminans* in iron-limited conditions, which benefits the multispecies biofilm community *in vitro* and promotes plant growth performance in iron-depleted soil. Our study reveals the molecular mechanisms underlying an emergent rhizosphere biofilm community function and demonstrates its importance in plant–microbe interactions.

## Introduction

Biofilms are complex aggregates of microorganisms attached to surfaces or to each other, composed of microbial cells embedded in extracellular polymeric substances. Biofilms are one of the primary modes of microbial existence in natural environments, including soil ^1,2^. This lifestyle provides numerous advantages to microorganisms, including improved nutrient acquisition, enhanced resistance to environmental stresses, and facilitation of horizontal gene transfer. In nature, biofilms typically contain multiple species; they have emergent community properties that cannot be predicted from the individual species alone and exceed the sum of contributions of individual member species of the community ^3,4^. Understanding and ultimately predicting emergent properties of biofilm communities remains a key challenge in microbial ecology ^5^. Although it is evident that emergent properties arise from interactions within the community, how these emergent features develop in biofilm communities and how they contribute to the advancement of biofilm populations or even influence their hosts remain poorly understood ^1,6^. For example, multispecies biofilms in the rhizosphere can provide plants with enhanced drought tolerance and pathogen suppression ^7,8^. These experimental observations suggest that rhizosphere biofilm communities can influence plant growth and health, yet a definitive link between biofilm community emergent properties and plant performance is yet to be established.

Metabolic cross-feeding is a widespread interaction in microbial communities and is particularly important in resource-limited environments ^9,10^. In multispecies biofilms, spatial proximity creates favorable conditions for metabolic cross-feeding, whereas the matrix-embedded structure minimizes diffusion losses and decreases the risk of metabolite exploitation by free-living microorganisms ^11^. Amino acid cross-feeding, particularly of branched-chain amino acids (BCAAs), is a common form of metabolic cross-feeding among bacteria ^12^. BCAAs not only serve as nutritional resources but also act as regulatory molecules of nonribosomal peptide synthesis ^13–15^. Other compounds, including vitamins, heme, organic acids, and sugars, have been implicated in microbial cross-feeding ^16–19^. Metabolite cross-feeding underpins the stability of biofilm communities. However, their rapid development and expansion may necessitate the management of other limiting resources.

Iron is an essential element for nearly all microbial life activities ^20^, participating in important processes such as electron transfer, DNA synthesis, and cellular respiration ^21,22^. However, in most natural environments, iron exits primarily as insoluble Fe (III) oxides with extremely low bioavailability ^23^. This limitation is particularly problematic in agricultural soils, especially in alkaline conditions, where iron deficiency significantly constrains both microbial activity and plant growth ^24^. To address this challenge, microorganisms produce and secrete high-affinity iron chelators called siderophores as a widespread, effective iron acquisition strategy ^25,26^. In the rhizosphere, microbial siderophores not only support bacterial iron acquisition but also improve plant iron nutrition ^27,28^. For example, bacillibactin, the siderophore produced by *Bacillus*, promotes plant iron acquisition in alkaline soil ^29^. Bacillibactin not only affects iron uptake but also influences the secretion of extracellular matrix and biofilm formation by *Bacillus* ^30^. For wild-type (WT) strains, the iron content in the extracellular matrix is 5-to-10-times higher than that inside the cells. In contrast, mutants with disrupted bacillibactin synthesis genes exhibit a significant decrease in extracellular matrix secretion capability and lose the ability to enrich iron in the extracellular matrix ^31,32^. Our latest research indicates that bacillibactin also acts as a signaling molecule that activates plant iron uptake, thereby enhancing iron absorption in iron-limited conditions ^29^.

Traditionally, siderophores have primarily been regarded as mediators of interspecies competition ^33^, where microorganisms compete for limited iron resources through the production of siderophores with different affinities, or by exploiting the siderophores produced by competitors. This competitive paradigm has been well-documented in pathogens, such as *Pseudomonas aeruginosa* and *Staphylococcus aureus*, which engage in fierce competition for limited iron resources ^34,35^. Similar competition occurs in soil, where siderophore-producing *Pseudomonas* can inhibit competitors through iron sequestration ^36^. Additionally, some microorganisms can directly use siderophores produced by other species, which has been observed in interactions between *P. aeruginosa* and *Mycobacterium* species ^37^. The interspecies exploitation of siderophores raises intriguing questions: Can siderophore sharing promote cooperation among microorganisms? Additionally, if siderophores function as a public good induced within biofilm communities, can these specialized metabolites also act as signals in the rhizosphere to modulate plant–microbe interactions?

This study explored the community-level emergent properties of a trispecies community of soil bacteria consisting of *Acinetobacter baumannii* XL380 (Ab), *Burkholderia contaminans* XL73 (Bc), and *Bacillus velezensis* SQR9 (Bv), that exhibits synergistic biofilm formation. We discovered a signaling and metabolite sharing network, including cross-feeding between Bv and the other two species through 5-aminovaleric acid (Bv to Bc/Ab) and BCAAs (Bc to Bv). Simultaneously, Bc secretes signaling molecules, including acyl-homoserine lactones (AHLs), that upregulate bacillibactin biosynthesis in Bv, which is exploited by Bc, explaining the enhanced biofilm community productivity. The synergistic interaction among the three species promotes plant growth performance in iron-deficient soil. We reveal a reciprocal mutualism that challenges traditional competitive interaction models in bacteria, and demonstrate how metabolic complementation can transform resource limitation into community-wide benefits.

## Results

### Synergistic biofilm formation by the trispecies community

We have previously collected bacterial isolates from cucumber rhizosphere soil and performed static cultivation, which demonstrated enrichment of Ab, Bc, and Bv in pellicle biofilms ^38,39^. Biomass quantification revealed that a trispecies biofilm formed by Ab, Bc, and Bv (AbBcBv) had significantly higher fresh weight than any mono- or dual-species biofilms (Fig 1A). Multispecies biofilms exhibited more robust and wrinkled morphology than single-species pellicle biofilms.

**Fig 1.**
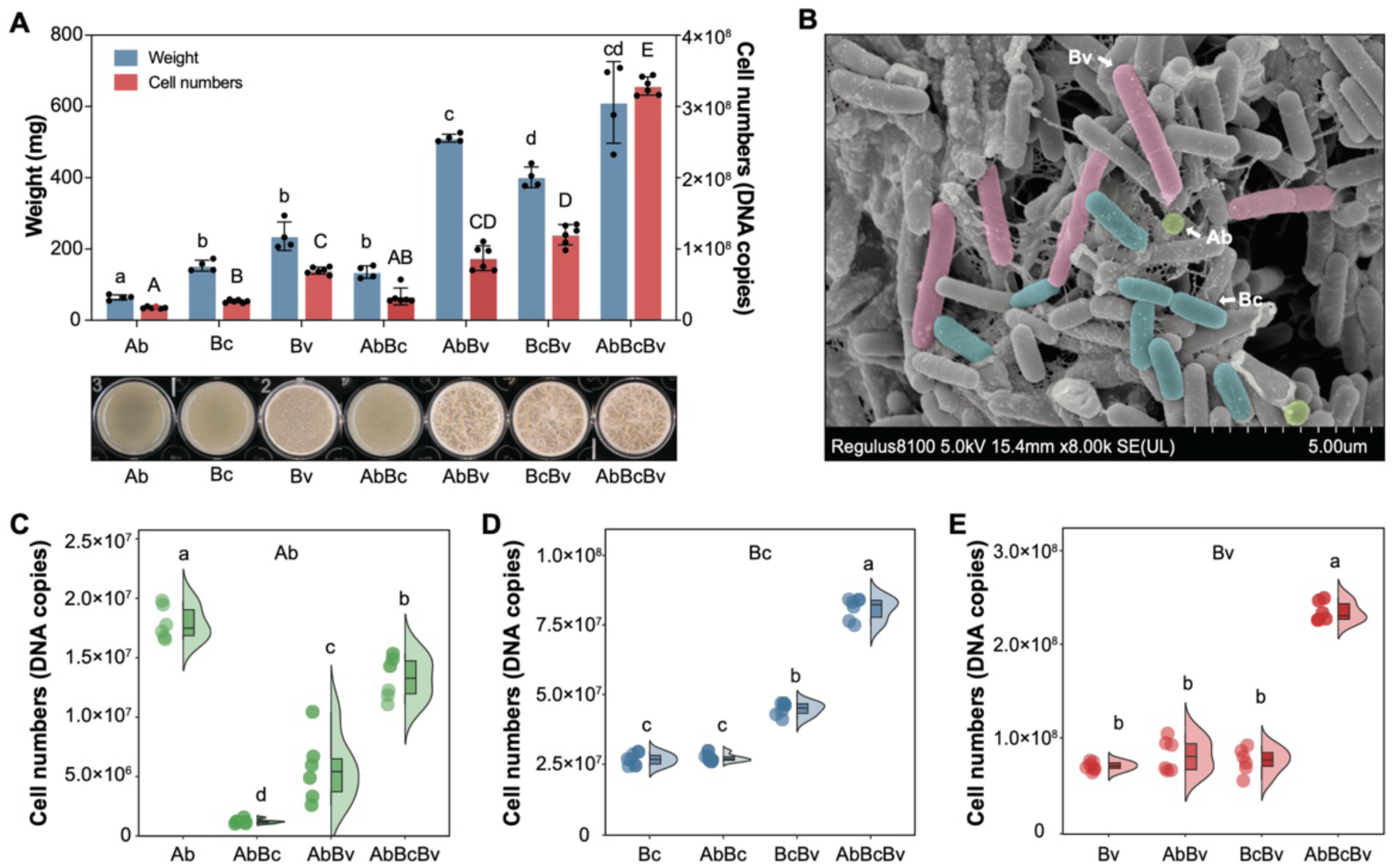
Synergistic biofilm formation. **(A)** Biofilm phenotype and biomass quantification. The well diameter was 15.6 mm. The left *y*-axis indicates the fresh weight of biofilms. Data presented are the mean ± s.d. (*n* = 4). The right *y*-axis displays the total cell numbers of biofilms as determined by quantitative PCR (qPCR). Data presented are the mean ± s.d. (*n* = 6). **(B)** Scanning electron microscopy of synergic biofilms. Green: *Acinetobacter baumannii* (Ab); blue: *Burkholderia contaminans* (Bc); red: *Bacillus velezensis* (Bv). **(C**–**E)** Cell numbers of Ab **(C)**, Bc **(D)**, and Bv **(E)** in different treatments. Data presented are the mean ± s.d. (*n* = 6). Different letters indicate statistically significant (*p* < 0.05) differences according to one-way analysis of variance (ANOVA) followed by Tukey’s honestly significant difference (HSD) *post-hoc* test.

To determine whether the enhanced biofilm in the coculture resulted from mutualism or competition, we quantified the cell numbers of each species. In the trispecies biofilm, Bv was the most abundant species (∼2.2×10^8^), followed by Bc (∼7.5×10^7^), and Ab (∼1.3×10^7^). The observation was confirmed by scanning electron microscopy (SEM), which showed a low abundance of Ab closely associated with abundant Bc and Bv (Fig 1B, Fig S1). The trispecies combination (AbBcBv) demonstrated the highest total cell number among all groups, significantly exceeding those in any other culture condition. The cell numbers of Bc (Fig 1D) and Bv (Fig 1E) were significantly higher than in their respective monocultures, whereas Ab decreased compared with its monoculture (Fig 1C). Pairwise comparisons revealed variable interaction patterns. Both AbBv and BcBv showed significantly higher cell numbers than their respective monocultures, whereas AbBc showed no significant difference compared with Bc alone (Fig 1A, right axis, *p* < 0.05). These findings suggest the enhanced trispecies biofilm results from complex interactions involving both mutualism and competition, where Bc and Bv benefit from the multispecies environment whereas Ab, though decreased in amount, remains an integral member of the community.

### Transcriptomic analysis of the trispecies biofilm

To elucidate the molecular details governing synergistic interactions, global gene expression profiles were systematically investigated. Compared with their monocultures, Bc and Bv differentially expressed >1000 and hundreds of genes, respectively, in BcBv dual-and AbBcBv trispecies biofilms (Fig 2A). Lists of the differentially expressed genes in BcBv and AbBcBv biofilms showed substantial overlap, including 1643 genes in Bc and 248 genes in Bv (Fig 2B,C). In contrast, only a few genes were differentially expressed in AbBc and AbBv combinations compared with monocultures (Fig 2A). Therefore, Bc and Bv are the primary interacting species that drive major transcriptomic changes, whereas Ab exerts limited effects on the transcriptional landscapes of the other community members.

**Fig 2.**
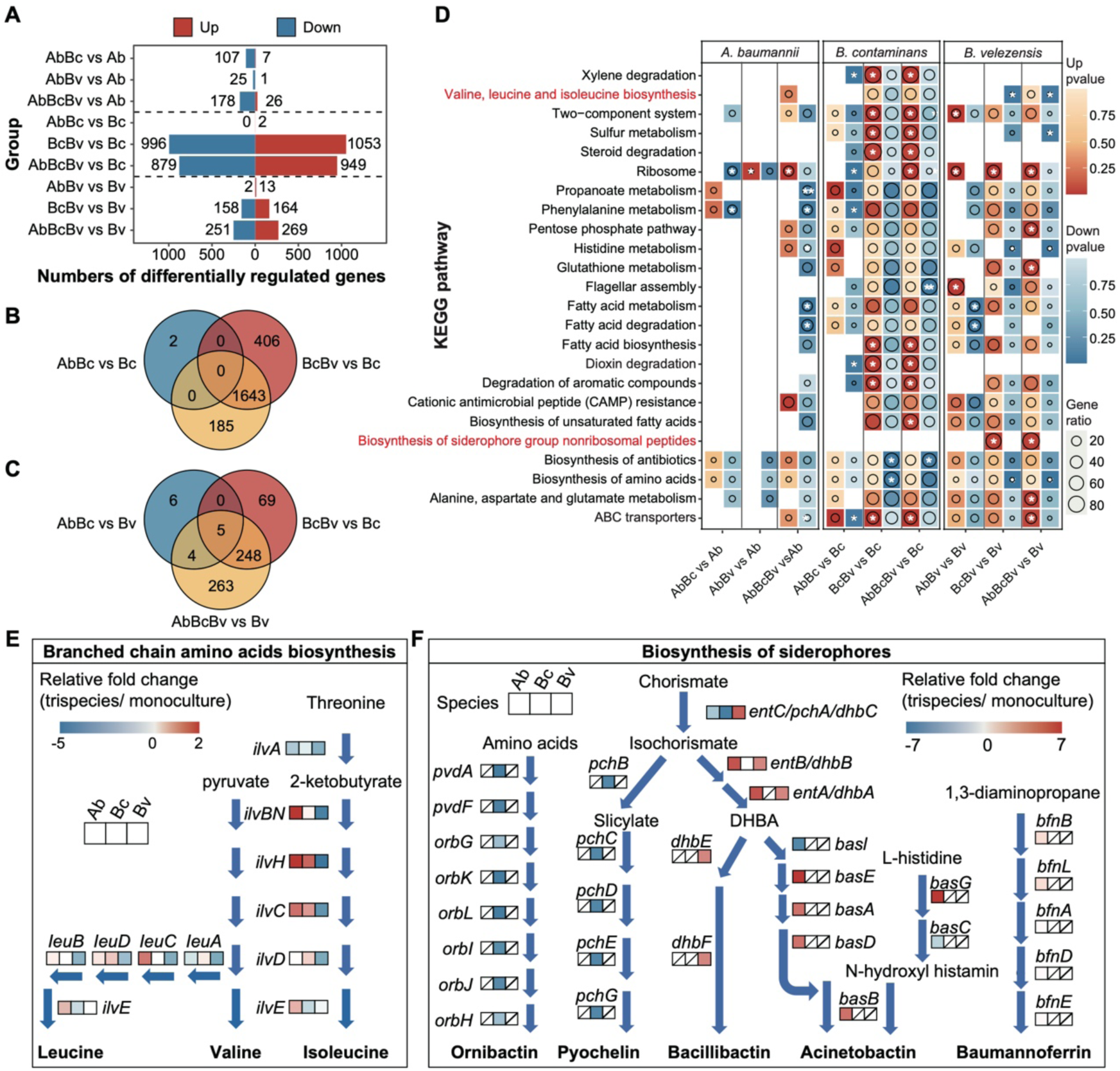
Transcriptome response in multispecies biofilms. **(A)** Numbers of differentially regulated genes in coculture compared with monoculture (log_2_ fold-change > 2, *p* < 0.05, *t*-test). The overlap of differentially upregulated **(B)** and downregulated **(C)** genes between different comparison groups. **(D)** Differentially regulated Kyoto Encyclopedia of Genes and Genomes (KEGG) pathways in multispecies biofilms compared with monoculture. An asterisk (*) indicates that a pathway was significantly regulated (*p* < 0.05, Fisher’s exact test, false discovery rate was adjusted by Benjamini–Hochberg correction). The pathway significance is color-coded, as shown (color represents the *p*-value). **(E** & **F)** Transcriptome results for relative gene expression in branched-chain amino acid (BCAA) biosynthesis pathways **(E)** and siderophore biosynthesis pathways **(F)** in trispecies coculture compared with monoculture. Because the siderophore biosynthesis pathways of Ab and Bc have not been incorporated into the KEGG database, they have been plotted manually.

Functional analysis of multispecies biofilms compared with monocultures revealed that 24 Kyoto Encyclopedia of Genes and Genomes (KEGG) pathways were differentially regulated (*p* < 0.05) (Fig 2D). Several KEGG pathways of Bc were significantly upregulated both in BcBv and the trispecies biofilm, including xylene degradation, two-component systems, sulfur metabolism, and ABC transporters. Conversely, genes encoding the biosynthesis of antimicrobials were significantly downregulated. In Bv, the gene cluster encoding siderophore biosynthesis was significantly upregulated in both BcBv and trispecies biofilms compared with monoculture. The genes encoding the ribosomes of Bv were significantly upregulated in all coculture groups, suggesting enhanced protein synthesis and faster growth, which is consistent with the higher cell number observed in mixed species biofilms. Conversely, genes encoding the biosynthesis of BCAAs (valine, leucine, and isoleucine) were significantly downregulated in BcBv and trispecies biofilms, suggesting decreased BCAAs production by Bv.

Given that Bv emerged as the predominant species in biofilm formation, we specifically focused on two pathways that were significantly regulated in multispecies biofilms compared with Bv monoculture: the upregulated siderophore pathway, and the downregulated BCAA synthesis pathway. Real-time quantitative -PCR (RTq-PCR) experiments confirmed that transcription of all genes for biosynthesis of bacillibactin (the siderophore of Bv) was upregulated in the triculture biofilm (Fig 2F, Fig S2A). In contrast, the transcription of all genes for biosynthesis of pyochelin and ornibactin (siderophores of Bc) was downregulated (Fig 2F, Fig S2B). Regarding the siderophores of Ab, 75% of the biosynthesis genes encoding acinetobactin synthesis were upregulated in the triculture biofilm, whereas the transcription of genes encoding the baoumannoferrin biosynthesis pathway remained unaffected. In the BCAA biosynthesis pathway, the *ilvC* and *ilvH* genes of Ab and Bc were upregulated in the trispecies biofilm but downregulated in Bv compared with monocultures (Fig 2E, Fig S2C). These expression patterns suggest that Bv may provide siderophore for the biofilm community, whereas Ab and Bc provide BCAAs in return, establishing a potential metabolic complementation relationship.

### Bc induces bacillibactin production in Bv and uses it for iron acquisition

To explore the importance of iron and siderophores in synergistic biofilm formation, we used three mutants of Bv ^30^: *Δdhb*, which is unable to produce bacillibactin and its precursor 2,3-dihydroxybenzoate (DHBA); *ΔdhbF*, which produces DHBA but not bacillibactin; and *ΔfeuB*, which lacks Fe–bacillibactin transport into the cells ^40^ (Fig S3). Using trispecies cocultures with either WT or mutant strains of Bv, we tested three different iron levels by adding extra FeCl_3_ or the iron chelator 2-dipyridyl (2DP) to standard tryptic soy broth (TSB) culture medium (Fig 3A, Fig S3). In low-iron (2DP-supplemented) medium, Co-*Δdhb*, Co-*ΔdhbF*, and Co-*ΔfeuB* exhibited severely impaired growth compared with Co-WT (Co-denotes coculture), including significantly decreased cell numbers (Fig 3A,B). This suggests that siderophore produced by Bv supported the growth of the biofilm community in iron-limited conditions. In high-iron (FeCl_3_-supplemented) medium, all cocultures showed similar biofilm phenotypes, because abundant iron was directly accessible without a requirement for siderophores (Fig 3A). In TSB (intermediate level of iron), the biofilms of Co-*Δdhb*, Co-*ΔdhbF*, and Co-*ΔfeuB* appeared thinner than Co-WT, and they displayed decreased cell numbers (Fig 3A,B), confirming the necessity of Bv siderophore in enhancing trispecies biofilm formation. Biofilms of Co-WT formed in high-iron conditions had significantly lower cell numbers than those in normal iron conditions, suggesting that siderophores play an additional role in the interspecies interaction, beyond iron scavenging.

**Fig 3.**
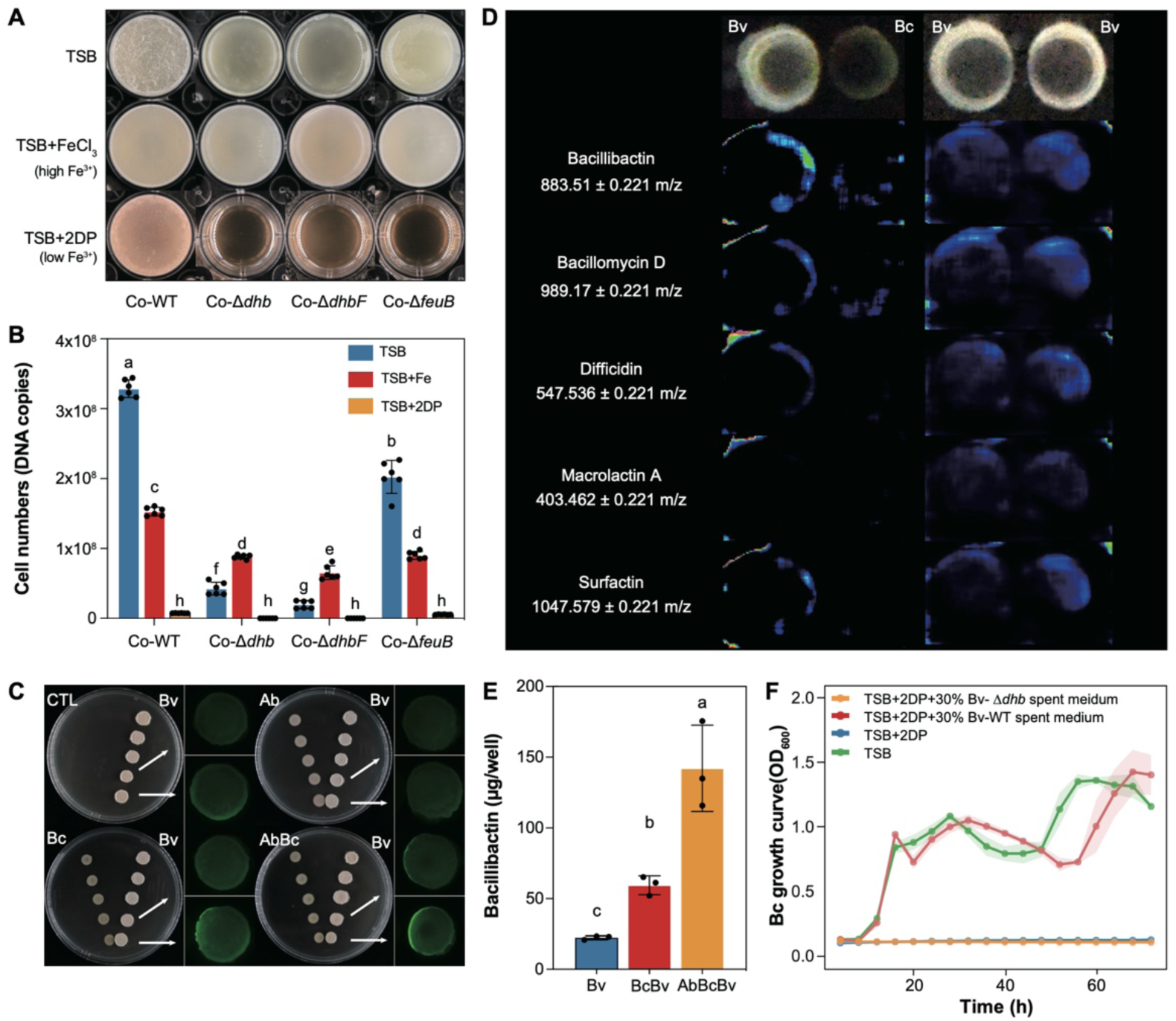
Enhanced bacillibactin production by Bv in interspecies interaction. **(A)** Phenotype of trispecies biofilms formed by Ab, Bc, and wild-type (WT) or mutants of Bv in tryptic soy broth (TSB) with three different iron concentrations. FeCl_3_ or 2-2‘-dipyridyl (2DP) was added at a final concentration of 0.5 mM to adjust the iron concentration. **(B)** Total cell numbers in the biofilms, determined by qPCR. Data presented are the mean ± s.d. (*n* = 6). **(C)** Induction of the *dhbABCEF* operon promoter in Bv upon contact with Bc. **(D)** Matrix-assisted laser desorption/ionization time-of-flight mass (MALDI-TOF) spectrometry imaging of bacillibactin during interactions. **(E)** The concentration of bacillibactin in biofilms. **(F)** Growth curves of Bc in different media. TSB with normal iron concentration was the control. TSB + 2DP was medium with a low iron concentration. The red line indicates medium containing 30% Bv spent culture supernatant, which contains bacillibactin. The yellow line indicates medium containing 30% spent culture supernatant of Bv mutant Δ*dhb*, which does not produce bacillibactin. Data presented are the mean ± s.d. (*n* = 5). Different letters indicate statistically significant differences (*p* < 0.05) according to one-way ANOVA followed by Tukey’s HSD *post-hoc* test.

Next, we monitored *dhbA* expression in WT Bv when it was cocultured with Ab or Bc. As the distance between colonies on an agar plate was decreased, Bc induced *dhbA* expression in Bv, whereas a comparable effect was not observed when Bv was paired with Ab (Fig 3C). The combination of Ab and Bc induced higher expression than did Bc alone, indicating that although Ab did not directly induce *dhbA* expression in Bv, it enhanced the interaction between Bc and Bv. To validate Bc-induced bacillibactin production in Bv, matrix-assisted laser desorption/ionization time-of-flight mass spectrometry (MALDI–TOF) imaging was used, which confirmed an increased level of bacillibactin in the Bv colony in the presence of Bc (Fig 3D). Whereas bacillibactin production was significantly enhanced, the spatial distribution of other Bv-produced secondary metabolites showed varying responses in the presence of Bc: bacillomycin D and surfactin showed moderate increases, whereas the polyketides difficidin and macrolactin A were not changed. Quantitative analysis of coculture biofilms revealed an elevated bacillibactin concentration in the trispecies biofilm (142.09 ± 17.59 ug/well) compared with Bv monoculture (22.44 ± 0.77 ug/well) (Fig 3E, Fig S4). These results confirmed that the presence of Bc increased bacillibactin production in Bv.

Next, we evaluated whether Bv siderophores can support Bc growth at different iron levels (Fig 3F). In low-iron medium (TSB + 2DP), Bc showed no growth when cultured alone or when supplemented with spent medium from culture of the Bv *Δdhb* strain. However, spent medium from culture of WT Bv restored the growth of Bc to a similar level to that in TSB without iron depletion, demonstrating that Bc can use bacillibactin produced by Bv to overcome iron limitation.

We then hypothesized that Bc secretes signaling molecules that induce siderophore production in Bv. Previous studies have demonstrated that AHLs produced by *Burkholderia cepacia* can regulate siderophore production ^41^. We thus tested induction of *dhbA* expression in Bv by eight AHL homologs (Fig S5). At 12 and 18 h, C4-HSL and C6-HSL significantly induced *dhbA* expression compared with the negative control, with C4-HSL showing induction levels comparable to that in the positive control (Bc supernatant) by 18 h (Fig S5A,B). High-performance liquid chromatography (HPLC) analysis confirmed the presence of C4-HSL in Bc supernatant (Fig S6).

In conclusion, a sophisticated chemical interplay is present within the biofilm community: Bc stimulates bacillibactin production in Bv, possibly through specific AHLs; Bv provides an iron-scavenging siderophore that supports Bc growth in iron-limited conditions; and Ab reinforces this mutualism, establishing a synergistic relationship critical for the enhanced trispecies biofilm formation.

### Ab and Bc provide BCAAs to support the growth of Bv

To validate the hypothesis that Bv exploits BCAAs produced by Ab and Bc in the mixed species biofilm, we cocultured Bv mutants Δ*ilvA*, Δ*ilvCH*, Δ*ilvD* (unable to synthesize all three BCAAs) and Δ*leuBCD* (unable to synthesize leucine)^42^ with Ab and Bc in TSB (Fig 4A). The biomass of the resulting coculture biofilms showed a clear increase compared with monoculture biofilms. Compared with the monocultures, the cell numbers of the WT, Δ*ilvA*, Δ*ilvD*, and Δ*ilvCH* strains exhibited a significant increase, whereas strain Δ*leuBCD* showed minimal growth enhancement (Fig 4B). These results demonstrated that Ab and Bc can provide BCAAs, particularly valine and isoleucine, to support the growth of Bv in the trispecies biofilm, confirming the existence of BCAA cross-feeding between these species.

**Fig 4.**
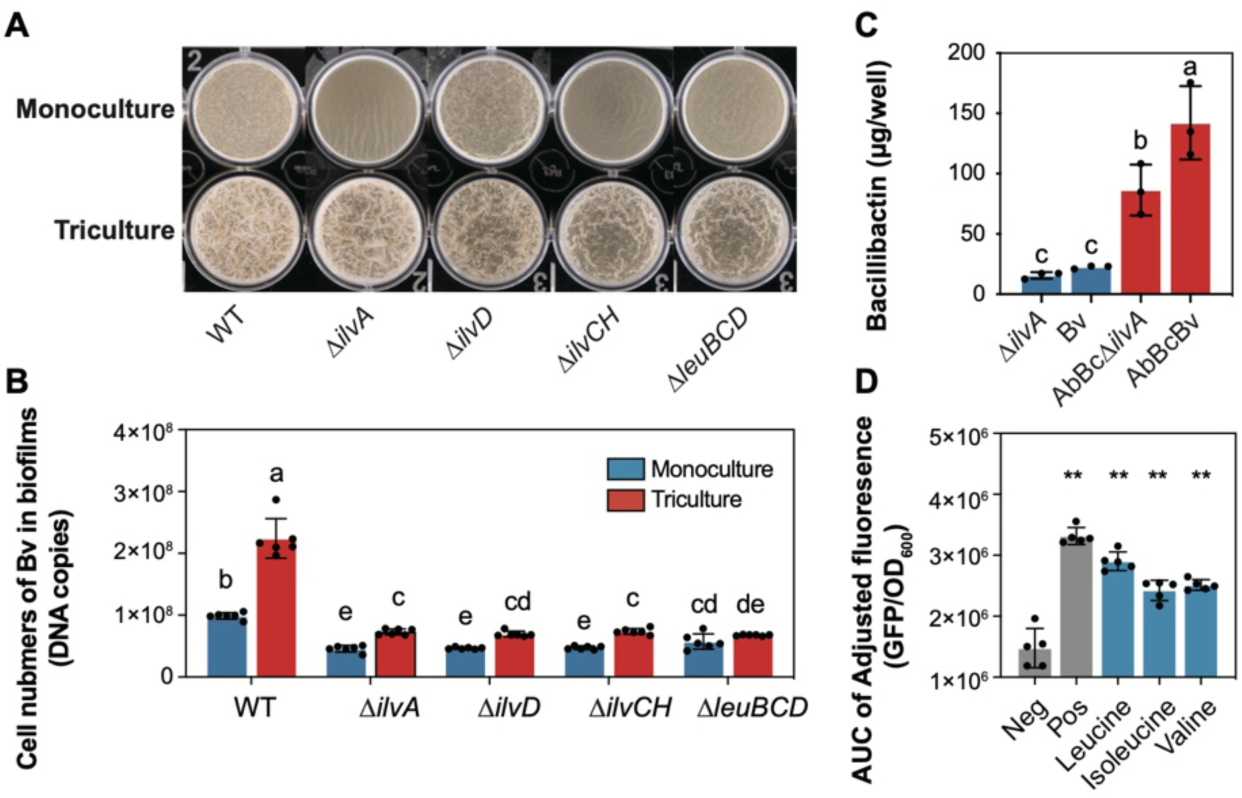
Triculture promoted the growth of Bv BCAA-deficient mutants in biofilms. **(A)** Phenotype of pellicles formed by BCAA-biosynthesis mutants of Bv in monoculture or in coculture with Ab and Bc. **(B)** Cell numbers of Bv in monoculture and coculture. Data presented are the mean ± s.d. (*n* = 6). **(C)** The concentration of bacillibactin in biofilms. Data presented are the mean ± s.d. (*n* = 3). Different letters indicate statistically significant (*p* < 0.05) differences according to one-way ANOVA followed by Tukey’s HSD *post-hoc* test. **(D)** Expression level of *dhb* of Bv in TSB supplemented with BCAAs as quantified by green-fluorescent protein (GFP) value (GFP/OD_600_). Data presented are the mean ± s.d. (*n* = 5). Statistical analysis was performed by *t*-test. (\**p* < 0.05, \*\**p* < 0.01).

BCAAs have been shown to influence secondary metabolite biosynthesis pathways ^43^, which is connected with nonribosomal peptide synthetases (NRPSs) ^44^. To examine whether BCAA availability affects bacillibactin production in Bv, bacillibactin levels were quantified (Fig 4C). In monocultures, the BCAA-deficient mutant Δ*ilvA* showed decreased bacillibactin production compared with the WT strain. Furthermore, direct supplementation of individual BCAAs significantly induced *dhbA* expression in Bv compared with negative controls, with the positive control (Bc spent medium) showing the highest level of induction (Fig 4D). When the BCAA-deficient mutant was cocultured with Ab and Bc in trispecies biofilms, bacillibactin production was significantly restored (Fig 4C). This restoration of bacillibactin production in the mutant coculture demonstrates that BCAAs secreted by Ab and Bc can be used by Bv to support siderophore biosynthesis, Bv growth, or both at the same time.

### Bv supplies metabolites that promote the growth of Bc and Ab

To explore other potential cross-feeding interactions in the trispecies community, the influence of spent medium on growth was tested (Fig 5A) ^42^. During cultivation in M9 medium, Bc depletes glucose, whereas residual glucose remains in Ac culture supernatant. The culture supernatant of Bc did not support growth of the other species (Fig S7). In contrast, spent medium from culture of Bv supported the growth of both Ab and Bc, indicating that Bv provides metabolites to the other two species.

**Fig 5.**
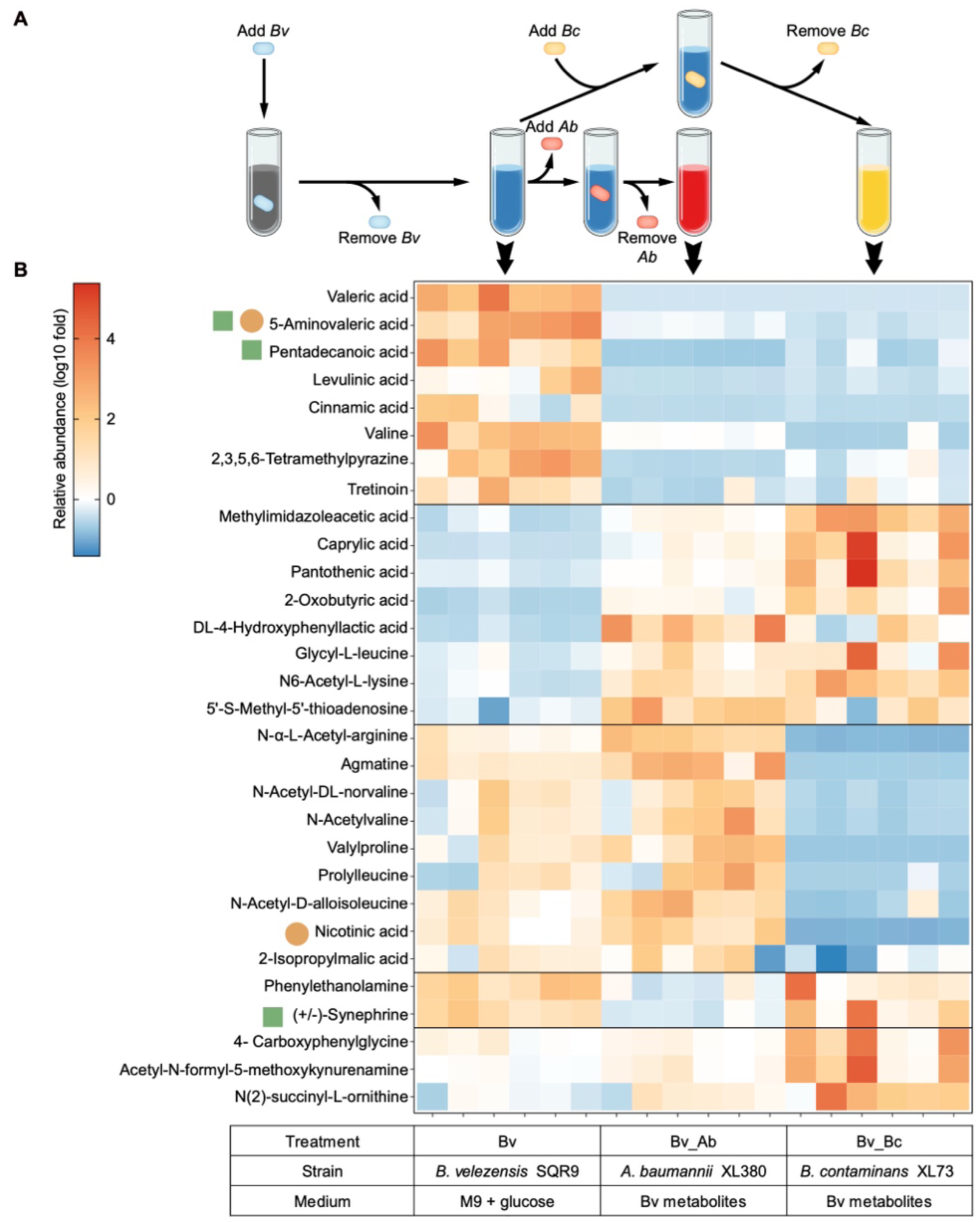
Metabolic facilitation stabilizes the trispecies cooperation. **(A)** Schematic representation of the spent medium assay. **(B)** Metabolic profiles of spent media. “Bv” indicates the spent medium of Bv grown on M9 glucose medium; “Bv_Ab” indicates the spent medium of Ab grown on spent medium of Bv; and “Bv_Bc” indicates the spent medium of Bc grown on spent medium of Bv. An orange cycle indicates a substance that can be used by Bc, as verified by growth assay. A green square indicates a substance that can be used by Ab, as verified by growth assay.

The top 30 differential compounds were analyzed based on the relative abundance (Fig 5B). Highly abundant compounds in Bv spent medium that were depleted after cultivation of Ab or Bc were designated as cross-fed compounds. These included valeric acid, 5-aminovaleric acid, pentadecanoic acid, levulinic acid, cinnamic acid, valine, 2,3,4,5-tetramethylpyrazine, and tretinoin. Additionally, species-specific use patterns were observed: nine compounds (*N*-α-L-acetyl-arginine, agmatine, *N*-acetyl-DL-norvaline, *N*-acetylvaline, valylproline, prolylleucine, *N*-acetyl-D-alloisoleucine, nicotinic acid, and 2-isopropylmalic acid) were used exclusively by Bc, whereas two compounds (phenylethanolamine and synephrine) were consumed specifically by Ab. Testing the impact of identified compounds on bacterial growth revealed that Ab grew on synephrine, 5-aminovaleric acid, and pentadecanoic acid, whereas Bc used 5-aminovaleric acid and nicotinic acid (Fig S8).

### Plant growth promotion in iron-deficient soil

To evaluate the ecological significance of iron-mediated interaction of the trispecies biofilm community, plant-growth-promoting (PGP) traits of the community were tested. The trispecies community demonstrated superior PGP traits, including enhanced phosphate solubilization (Fig 6A,B), ammonia production (Fig 6C,D), siderophore production (Fig 6E), and exopolysaccharide production (Fig 6F), compared with mono- and dual-species treatments. Growth of *Arabidopsis thaliana* was severely stunted in the iron-deficient soil without inoculation (the control, CTL), confirming the detrimental influence of iron limitation on plant growth (Fig 6G,H). Plants inoculated with the trispecies consortium (AbBcBv) exhibited the most robust growth and highest fresh weight, significantly outperforming all other treatments (Fig 6G,H). Critically, when the bacillibactin-deficient mutant (Bv-Δ*dhb*) was used instead of WT Bv, plant growth was substantially decreased, demonstrating the critical role of bacillibactin in plant iron nutrition in iron-deficient conditions. Dual-species combinations showed intermediate growth promotion effects, with BcBv performing better than AbBv and AbBc. This plant growth promotion pattern correlated with the biofilm productivity, where BcBv demonstrated stronger synergistic interactions than the other dual-species combinations.

**Fig 6.**
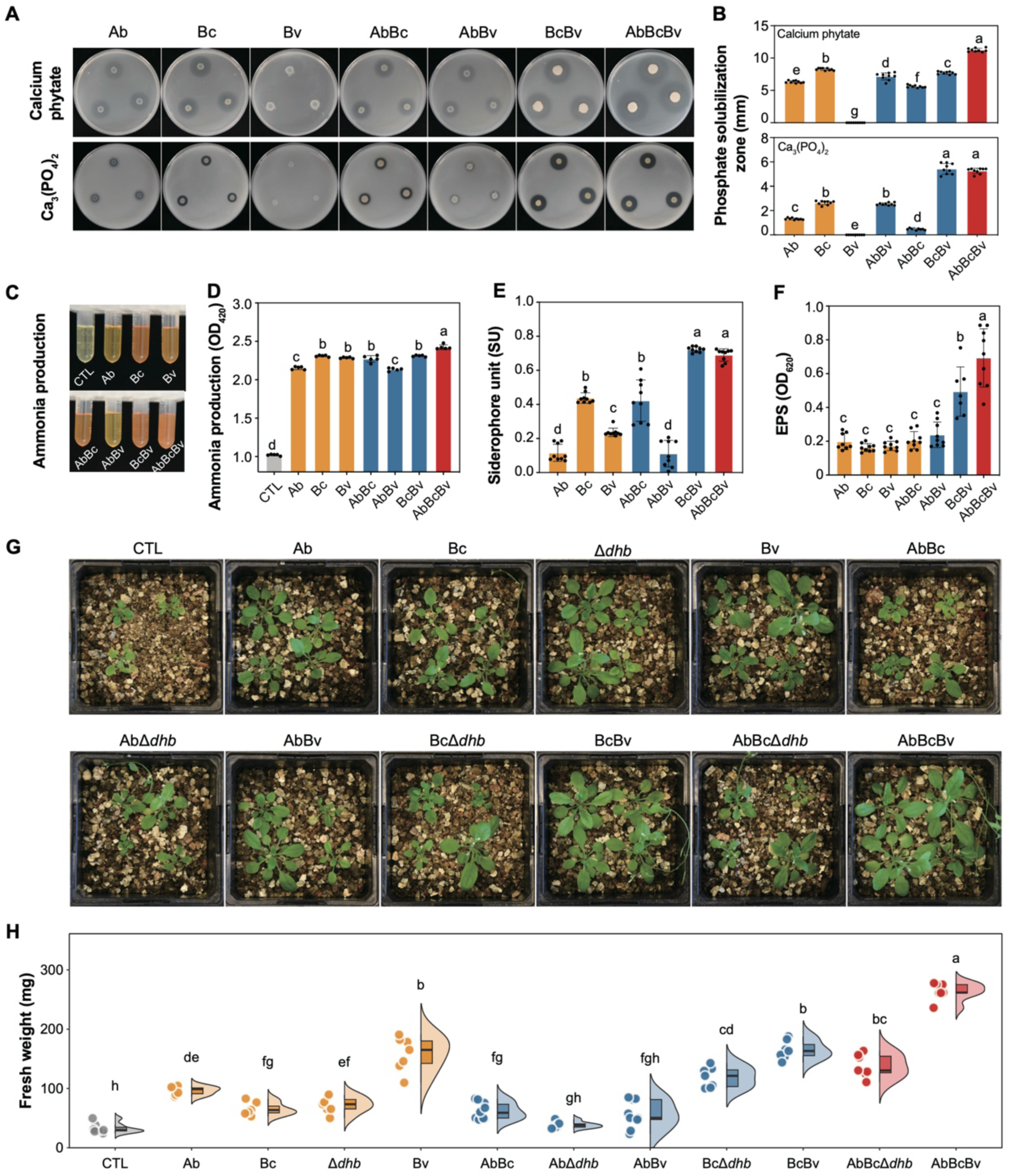
Multispecies community enhanced plant growth in low-iron conditions. The phosphate [Ca_3_(PO_4_)_2_ and calcium phytate] solubilization **(A** & **B)**, ammonia production **(C** & **D)**, siderophore production **(E)**, and exopolysaccharide production **(F)** in different treatments. The plate diameter was 9 cm. Data presented are the mean ± s.d. (*n* = 5–9). **(G)** The growth of *Arabidopsis thaliana* in an iron-deficient environment with different microbial treatments. **(H)** The fresh weight of *A. thaliana*. Data presented are the mean ± s.d. (*n* = 8). Different letters indicate statistically significant (*p* < 0.05) differences according to one-way ANOVA followed by Tukey’s HSD *post-hoc* test. CTL: control, not inoculated with bacteria; Bv-Δ*dhb*: a mutant of Bv that cannot produce bacillibactin.

## Discussion

Here, we reveal emergent properties of a trispecies bacterial community, including enhanced biofilm formation, metabolic cross-feeding, elevated siderophore production, and improved plant growth promotion (Fig 7). Within the community, Bv serves as the primary metabolic supplier of essential metabolites, such as 5-aminovaleric acid, pentadecanoic acid, nicotinic acid, and synephrine, that support the growth of Bc and Ab, whereas Bc and Ab reciprocally deliver BCAAs. Biofilm synergism is further promoted by elevated bacillibactin production by Bv that promotes iron acquisition by the entire community; the increased bacillibactin production is stimulated both by the signaling molecule C4-HSL from Bc and by BCAAs from Bc and Ab. In iron-deficient conditions, this metabolic complementation not only secures community survival but also confers plant growth promotion effects. Our findings demonstrate that siderophores facilitate interspecies synergism in biofilms and enhance plant iron nutrition.

**Fig 7.**
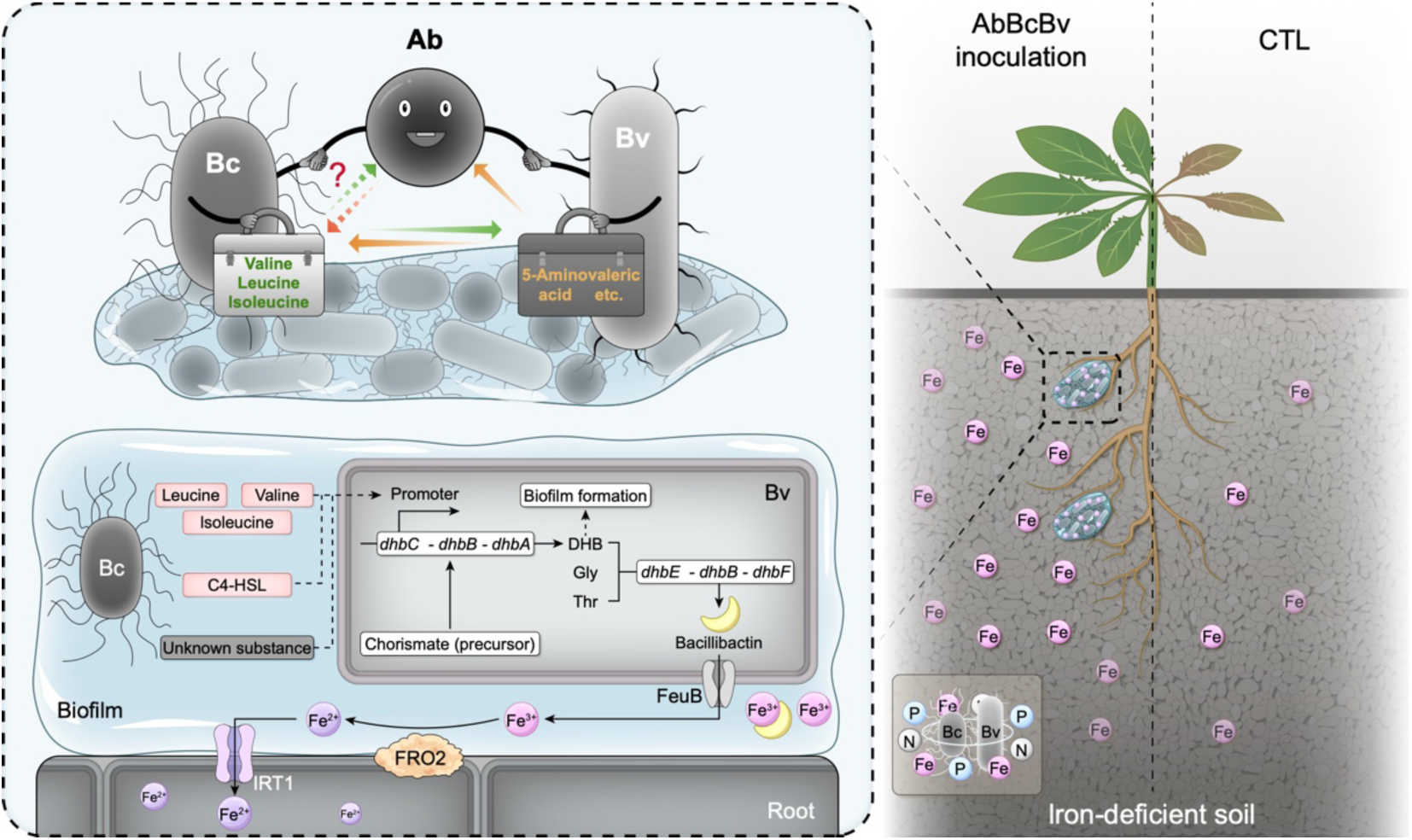
Summary figure. **(A)** Metabolic cross-feeding and iron mobilization mechanism. Mutual metabolic exchange occurs between Bc and Bv. Bc metabolites, including BCAAs (valine, leucine, isoleucine), C4-HSL, and other substances, induced and improved bacillibactin biosynthesis in Bv. Bacillibactin chelates Fe^3+^, and FeuB transports iron complexes. FRO2 reduces Fe^3+^ to Fe^2+^ for plant uptake via IRT1 in *Arabidopsis* roots. **(B)** AbBcBv consortium inoculation enhanced plant uptake of iron, ammonia and phosphate, promoting the growth of *A. thaliana* in iron-deficient soil.

The enhancement of the trispecies biofilm observed in this study represents emergent properties arising from complex interspecies interactions. The significantly higher biomass and cell numbers in AbBcBv treatment compared with monoculture demonstrate such emergent benefits, where the community performance surpasses what would be expected from simply combining the capabilities of individual species. Emergent properties in diverse multispecies biofilms are well-documented, including enhanced pathogen suppression by skin bacterial symbionts and improved stress tolerance in soil microbial communities ^45^. Skin bacterial symbionts with greater species richness exhibit superior suppression of the amphibian fungal pathogen *Batrachochytrium dendrobatidis* through combined dominant effects and complementarity mechanisms. Additionally, probiotic bacterial consortia consisting of up to eight *Pseudomonas* strains demonstrate emergent consortium-level effects in the tomato rhizosphere, where four- and eight-strain consortia reached densities up to 10-times higher than single-strain inoculants, exhibiting transgressive overyielding that could not be predicted from individual strain performance ^46^. These examples demonstrate how community-level emergent properties can provide ecological advantages that individual species cannot achieve alone. These emergent characteristics reflect a complex interplay of spatial organization, metabolic interdependencies, and chemical communication networks within structured microbial communities.

Metabolic cross-feeding is a fundamental mechanism that drives microbial community assembly and stability. This phenomenon has been extensively documented across diverse systems, from vitamin interdependencies ^17^ to amino acid exchange networks in synthetic communities ^47^. Metabolic interactions can augment biofilm formation in dual-species systems (e.g., heme cross-feeding in *Staphylococcus aureus* and *Enterococcus faecalis* biofilms), and multigenome metabolic modeling predicted functional interdependencies in plant microbiomes ^18,48^. Among the three dual-species treatments tested here, BcBv exhibited the strongest synergistic effects, a pattern consistently validated across transcriptomic and metabolomic analyses. Metabolome analysis revealed a biochemical basis underlying the mutualism between Bc and Bv, where Bv functions as the primary metabolite provider, secreting compounds including 5-aminovaleric acid, pentadecanoic acid, nicotinic acid, and synephrine that support community growth. Reciprocally, Ab and Bc provide BCAAs that promote the metabolic processes of Bv, particularly siderophore biosynthesis. BCAA cross-feeding has also been reported in other PGP rhizosphere biofilm communities ^7,42^ suggesting that amino acid cross-feeding represents a widespread strategy in multispecies biofilms. γ-Aminobutyric acid (GABA) can function as a cross-kingdom signal, mediating communication between bacteria, influencing gene expression and behaviors, and thereby modulating the composition and dynamics of microbial communities ^49^. We revealed that a Bv-secreted GABA homolog, 5-aminovaleric acid, supports the growth of Ab and Bc; however, its potential signaling function remains to be explored.

In natural environments, iron limitation represents one of the primary challenges faced by microorganisms, particularly in alkaline soils and rhizosphere environments ^16^. Our study reveals that siderophore sharing can facilitate interspecies cooperation in biofilm communities, contrasting with the conventional view of siderophores primarily being competitive tools. Such cooperative siderophore-mediated community performance challenges the traditional competition-focused models and suggests a mutualistic framework where iron acquisition becomes a community-wide benefit rather than an individual advantage. The ecological relevance of this siderophore-mediated cooperation was validated through PGP experiments, where the trispecies community significantly promoted the growth of *Arabidopsis* in iron-deficient soil. The use of bacillibactin-deficient mutants confirmed that cooperative siderophore production was essential for optimal PGP, directly linking laboratory observations of microbial cooperation to a tangible ecological outcome. These findings demonstrate how interspecies cooperation in biofilm communities can enhance both microbial survival and plant performance. The siderophore-mediated interaction extends beyond microbial communities to encompass cross-kingdom interactions, where bacterial siderophores can directly facilitate plant iron acquisition, ultimately benefiting both microbial survival and plant health in iron-limited conditions ^28,29^.

The regulatory network underlying the trispecies biofilm represents a highly integrated metabolic system where signaling molecules and nutritional exchanges contribute to community function. The simultaneous upregulation of siderophore biosynthesis and downregulation of BCAA synthesis pathways in Bv suggests a complementary expression pattern that demonstrates metabolic coupling between amino acid metabolism and siderophore synthesis. From a biosynthetic perspective, bacillibactin synthesis requires a substantial of amino acid precursors and energy investment, particularly because its backbone structure formation depends on amino acid-derived building blocks ^50,51^. The downregulation of BCAA synthesis pathways accompanied by upregulation of protein synthesis-related genes suggests that Bv optimizes siderophore production through reallocation of amino acid resources ^20^. Formation of the trispecies biofilm induces numerous unique gene expression changes that could not be predicted from simple dual-species interactions, suggesting the existence of higher-order regulatory mechanisms in multispecies environments, such as cross-regulation of quorum sensing signals and system-level redistribution of metabolic flux ^52,53^. The AHL-mediated interspecies signaling mechanism exemplifies the sophisticated regulatory characteristics of this synergistic system, where Bc-derived C4-HSL and C6-HSL induce expression of the *dhbA* gene of Bv. Moreover, although Ab does not directly induce *dhbA* expression in Bv, it enhances the impact of Bc on Bv, suggesting that Ab may exert regulatory control by modulating signal molecule stability or the sensitivity of Bv to these interspecies signals. The decreased bacillibactin production in Bv in the absence of BCAAs further confirms the biochemical coupling between amino acid metabolism and siderophore synthesis, where BCAAs influence siderophore synthesis through multiple pathways: serving as direct substrates for peptide synthesis ^54^, functioning as energy sources to supply ATP and reducing power ^15^, and acting as metabolic regulatory signaling molecules ^13^. These integrated mechanisms create a mutualistic cycle that enhances biofilm formation, increases community biomass, and promotes plant growth in iron-deficient conditions.

Beyond the rhizosphere environment, internal metabolic patterns and emergent properties of multispecies biofilm communities are relevant in microbial coexistence across diverse habitats, drug resistance, and the spread of antibiotic resistance genes. Potential human pathogens often reside within multispecies biofilm communities. Therefore, a deeper understanding of the metabolic interactions and key signaling pathways within multispecies biofilms could motivate strategies for efficient disruption of biofilm communities and precise elimination of pathogenic bacteria. Future research could explore the influence of quorum sensing signals (such as AHLs) on siderophore production in microbiomes, as well as the universality of cross-kingdom signal regulation in multispecies biofilm communities.

## Materials and Methods

### Strains and growth conditions

Bacterial strains and plasmids used in this study are listed in Table S1. *Bacillus velezensis* SQR9 (China General Microbiology Culture Collection Center, CGMCC no. 5808, NCBI accession no. PRJNA227504), *Acinetobacter baumannii* XL380 (Agricultural Culture Collection of China, ACCC61689, NCBI accession no. PRJNA593376), and *Burkholderia contaminans* XL73 (ACCC61690, NCBI accession no. PRJNA593683) were isolated from the cucumber rhizosphere ^38^. Starting inoculum was prepared from overnight cultures grown in TSB (Hopebio HB4114; 30°C, 180 rpm), harvested by centrifugation (6,000 x g for 2 min), and resuspended in 0.9% NaCl solution to achieve an optical density at 600 nm of 1.0 (OD_600_ = 1.0). When required, medium was supplemented with zeocin (20 μg L^−1^) or chloramphenicol (5 μg L^−1^).

M9 medium was prepared using minimal salts (5×, Sigma-Aldrich, M6030, Germany) supplemented with 2 mmol L^−1^ MgSO_4_ and 0.1 mmol L^−1^ CaCl_2_. This basal medium was further amended with various sole carbon sources as detailed in Table S2. Water-soluble compounds were added at 10 mg mL^−1^, whereas compounds with limited solubility were added to their maximum solubility.

### Biofilm formation and quantification

For visualization, colony biofilms were formed by spotting 2 μL of starting inoculum onto solid TSB (1.5% agar) and incubating statically at 30°C for 24 h. Pellicle biofilms were cultivated in 24-well plates by inoculating 20 μL of the starting inoculum into 2 mL of liquid TSB and incubating at 30°C for 24 h. For coculture biofilms, equal volumes of Ab, Bc, and Bv (set to same OD_600_) starting inoculum were combined. Iron availability was modulated by supplementing TSB with either 0.1% 2DP stock solution (Macklin, B807242, China; 78.10 mg mL^−1^) to establish iron-restricted conditions, or 0.1% FeCl_3_·6H_2_O stock solution (131.6 mg mL^−1^) to establish iron-rich conditions.

Pellicle biofilm biomass was determined gravimetrically as described before ^42^. Briefly, 100 μL of starting inoculum was added into 10 mL of TSB in six-well plates containing sterile nylon mesh cell strainers (100-μm pore size). After 24 h of static incubation at 30°C, the strainers were taken out, excess liquid was removed, and the total weight was recorded. Biomass was calculated as the difference between the total weight and the weight of the strainer.

For strain-specific quantification of biofilm-associated cells, 100 μL of inoculum was added into 10 mL of TSB in six-well plates equipped with nylon mesh strainers containing sterilized 1.5-cm^2^ woven filters. After 24 h of static incubation at 30°C, filters carrying pellicle biofilms were transferred to 1.5-mL tubes and stored at −80°C for DNA extraction. For planktonic growth analysis, 40 μL of inoculum was added to 4 mL of TSB in sterile tubes and cultured at 30°C, 180 rpm, for 24 h, followed by centrifugation and storage at −80°C. Strain-specific primers were designed through comparative genomic analysis, as described before^42^ (see Table S3 for primer sequences and standard curves). Absolute quantification was performed in 20-μL reactions with ChamQ SYBR qPCR Master Mix (Vazyme, Q711, China) according to the manufacturer’s instructions. The thermal cycling conditions were: 95°C for 30 s, followed by 40 cycles of 95°C for 5 s and 60°C for 45 s, with standard melt curve analysis. All treatments included six biological replicates.

### Whole-genome transcriptomic analysis and qRT–PCR

Pellicle biofilms were harvested from woven filters as described above. RNA was extracted using the E.Z.N.A. Bacterial RNA Kit (Omega Bio-tek, USA). For transcriptomics, sequencing libraries were constructed with the NEBNext Ultra Directional RNA Library Prep Kit and sequenced on an Illumina HiSeq 2000 platform. Raw data have been deposited in the NCBI SRA database (BioProject accession number PRJNA1281721). Following quality filtration, reads were aligned to reference genomes using Bowtie2. Differential expression was analyzed with DESeq2 using false discovery rate (FDR) correction. Genes with log_2_ fold-change > 2 and FDR < 0.05 were considered significantly differentially regulated. Functional categorization and pathway enrichment were performed with EggNOG-mapper v2 for KEGG Orthology terms.

For qRT–PCR, RNA was reverse-transcribed into cDNA using the PrimeScript RT Reagent Kit. Expression profiles of selected genes [*dhbA*, *dhbF*, *btr*, *feuB*, *tasA*, and *epsD* of Bv; *pchA*, *pchG*, *pchF*, *pchE*, and *Ipr1* of Bc; as well as *ilvC* and *ilvH* of all three species (Ab, Bc, and Bv)] were determined by qRT–PCR using ChamQ SYBR qPCR Master Mix. The *recA* gene served as an internal control for Bv and Bc, whereas *gyrB* was used for Ab. Primer sequences are provided in Table S3. The PCR conditions were: 95°C for 10 s, followed by 40 cycles of 95°C for 5 s and 60°C for 34 s. Relative expression levels were calculated using the 2^−ΔΔCT^ method.

### Microscopy

Bacterial cell morphology was examined using transmission electron microscopy. Overnight cultures were harvested, washed with phosphate-buffered saline, adjusted to OD_600_ = 0.5, applied to formvar/carbon-coated copper grids, and negatively stained with 2% (w/v) uranyl acetate. Imaging was on a Hitachi HT7800 TEM (Hitachi High-Tech, Japan)

Biofilm architecture was analyzed by SEM. Samples were fixed in 2.5% glutaraldehyde and 2% osmium tetroxide, dehydrated through graded ethanol and *tert*-butanol, and sputter-coated, before imaging on an SEM Regulus 8100 (Hitachi High-Tech).

Fluorescent reporter expression in colony and pellicle biofilms was visualized with an Axio Zoom V16 stereomicroscope (Carl Zeiss, Jena, Germany). Green-fluorescent protein (GFP) signals were captured using excitation at 470/440 nm and emission at 525/550 nm. Exposure time was 50–100 ms with 15% LED intensity, and acquisition parameters were kept constant across all samples to allow comparison.

### Fluorescently labeled Bv strain construction

Fluorescent transcriptional reporters were created as described by Xu *et al*. ^8^ Briefly, promoter regions of target genes were fused to a *gfp* coding sequence via overlap PCR and cloned into vector pNW33n. The resulting constructs (primer sequences given in Table S3) were introduced into WT Bv and derived mutants.

### Growth curve assays

Bacterial planktonic growth was monitored in 96-well plates using an Agilent Synergy H1 microplate reader (BioTek, USA). Cultures were initiated by putting 2 μL of the starting inoculum into 200 μL of TSB or M9 medium, then incubated at 30°C with continuous shaking. OD_600_ was measured every 30 min for up to 96 h. For carbon source use tests, M9 medium was supplemented with individual sole carbon sources (Table S2).

For BCAA induction experiments, TSB was supplemented with leucine, isoleucine, valine, or a mixture (100 μg mL^−1^). Both OD_600_ and GFP fluorescence values (excitation/emission, 485/528 nm) were recorded every 30 min for 48 h. All treatments included five replicates.

### MALDI–TOF imaging mass spectrometry (MS)

Colony biofilms were incubated as described above. Spatial distribution of metabolites within colonies was analyzed using MALDI–TOF MS. Colonies were excised, transferred to glass slides, desiccated at 30°C for 2 h, and coated with 20 mg mL^−1^ 2,5-dihydroxybenzoic acid containing 1.0% trifluoroacetic acid. The prepared slides were analyzed on an UltraFlextreme MALDI TOF/TOF system (Bruker Daltonics, USA) in reflector negative mode, *m/z* 100–2000. Data were processed with SCiLS Lab 2014b software with total ion count normalization. Target secondary metabolites (bacillibactin, bacillomycin D, difficidin, macrolactin A, and surfactin in Bv) were analyzed based on specific *m/z* values.

### HPLC/MS analysis

Standard bacillibactin was obtained from Biophore Research Products (Universität Tübingen, Germany). Biofilm extracellular extracts were prepared by sonication of pellicles in water, followed by centrifugation and ethyl acetate extraction. Dried extracts were reconstituted in methanol and analyzed using an Agilent 6410B Triple Quadrupole LC/MS instrument coupled to an Agilent 1200 HPLC system, equipped with a reverse-phase column (XBridge C18 5 μm, 4.6 × 250 mm). The samples were resolved at 220 nm and 30°C, at a flow-rate of 0.3 mL min^−1^, with a linear gradient from 70% to 100% (0–20 min) acetonitrile/water solvent (0.1% formic acid), and at 70% acetonitrile from 20–30 min ^55^. Mass spectra were collected in negative electrospray ionization mode.

For AHL analysis, Bc culture supernatants were extracted with ethyl acetate, dried, and dissolved in methanol. Samples were analyzed on a Waters e2695 equipped with an XBridge C18 column (5 μm, 4.6 × 250 mm) with isocratic elution (methanol/water 65:35, 1 mL min^−1^, 30°C, 40 min).

### Cross-feeding assay and metabolome analysis

Individual strains were cultured in 100 mL of M9 medium with glucose as the sole carbon source at 30°C and 180 rpm, until glucose depletion by Bc and Bv. However, Ab could not consume all the glucose. Spent media were obtained by centrifugation and filtration, and used as growth substrates for reciprocal cultivation by inoculating 1% (v/v) of each strain into spent media from the other species. Growth was monitored by OD_600_ for up to 72 h.

Extracellular metabolites were analyzed by untargeted metabolomics using a Vanquish UHPLC system coupled to an Orbitrap Q Exactive HF-X mass spectrometer (Thermo Fisher Scientific, USA). Samples were separated on a Hyperil Gold C18 column (100 × 2.1 mm, 1.9 μm) with a 16-min linear gradient of water and methanol containing either 0.1% formic acid (positive mode) or 5 mmol L^−1^ ammonium acetate, pH 9.0 (negative mode). The flow-rate was 0.2 mL min^−1^ at 30°C. Mass spectra were acquired in both positive and negative electrospray ionization modes. Raw data were processed with Compound Discoverer 3.0 (Thermo Fisher Scientific) for peak alignment, feature extraction, and intensity normalization. Metabolites were annotated by matching to the mzCloud and ChemSpider databases, and further analyzed on the Majorbio Cloud Platform. Features with relative standard deviation >30% in quality control samples were excluded. Differential metabolites were identified by orthogonal partial least squares discriminant analysis combined with Student’s *t*-test (VIP ≥ 1, p < 0.05). Heatmaps of the top 30 differential metabolites were generated. Four biological replicates were used for each treatment.

### Analysis of PGP traits

Phosphate solubilization activity was examined on specific agar medium supplemented with either calcium phytate or Ca_3_(PO_4_)_2_ as insoluble phosphate sources, following an established protocol ^56^. Exopolysaccharide content within biofilm was determined using the phenol–sulfuric acid colorimetric assay ^57^. Ammonia production was evaluated by assay with Nessler’s reagent in peptone medium ^58^. Siderophore production was quantified using the Chrome Azurol S colorimetric assay with cell-free supernatants from cultures grown in modified King’s B medium ^33^. All assays were performed with three to six biological replicates, and absorbance was measured at appropriate wavelengths according to standard methods.

### Plant pot experiment design

*Arabidopsis thaliana* (Col0) seeds were surface-sterilized and germinated on Murashige and Skoog medium with 1% agar (Hopebio, HB8469). Seven-day-old uniform seedlings were transplanted into pots containing 250 g of a sterilized growth substrate composed of vermiculite and perlite (5:1, v/v). Four seedlings were planted per pot, with two replicates per treatment. The starting inoculum was mixed into the substrate to achieve a final density of 10^7^ colony-forming units g^−1^ substrate. Iron deficiency was induced by adjusting ¼ Murashige and Skoog liquid medium to pH 8.0 with KOH before watering. Plants were grown with a long-day photoperiod (16-h light/8-h dark) at 22°C and watered every 4 days. After 4 weeks, plant weight was measured. Uninoculated plants were set as the control. The inoculation treatments were as follows: Ab, Bc, Bv-*Δdhb*, Bv, AbBc, Ab-*Δdhb*, AbBv, Bc-*Δdhb*, BcBv, AbBc-*Δdhb*, and AbBcBv.

### Statistical analysis

Statistical analyses were performed using R version 4.4.1 or GraphPad Prism 10. Figures were generated using the ggplot2 R package, GraphPad Prism 10, and Adobe Illustrator 2025. Details of specific statistical tests, significance thresholds, and sample sizes are provided in the figure captions.

## Data availability

Genome data for *B. velezensis* SQR9, *A. baumannii* XL380 and *B. contaminans* XL73 are available under NABI BioProject accession number PRJNA227504, PRJNA593376 and PRJNA593683, respectively. Transcriptome raw data have been deposited in the NCBI SRA database (BioProject accession number PRJNA1281721). All other data generated and analyzed during this study are either available in figure legends or can be requested from the corresponding author.

## Acknowledgments

This work was financially supported by Jiangsu Provincial Key Basic Research Projects (BK20253034), and National Natural Science Foundation of China (32361143785, 42477310 and 32400113). JX was supported by a China Scholarship Council fellowship during his stay in Leiden. ÁTK was funded by the European Union (ERC, MicroClock, 101166968). Views and opinions expressed are however those of the author(s) only and do not necessarily reflect those of the European Union or the European Research Council Executive Agency. Neither the European Union nor the granting authority can be held responsible for them. MLS is supported by the Danish National Research Foundation (DNRF137).

## Author contributions

J.X., X.S., Á.T.K., Z.X., and R.Z conceived the project. J.X., X.S., K.D., and H.Z. performed the experiments. V.H.T and M.L. performed transcriptome analysis. W.X., N.Z., Á.T.K., Q.S., and R.Z. contributed to experimental design and methodology. X.S. performed the metabolomic analysis. J.X., K.D., and H.Z. conducted biofilm formation assays and microscopy analysis. J.X. performed gene knockout experiments, MALDI-TOF imaging and plant growth experiments. J.X., X.S., W.X., and N.Z. performed data analysis and statistics. J.X., X.S., and Á.T.K wrote the manuscript and corrections from all authors.

---

**FIG S1.**
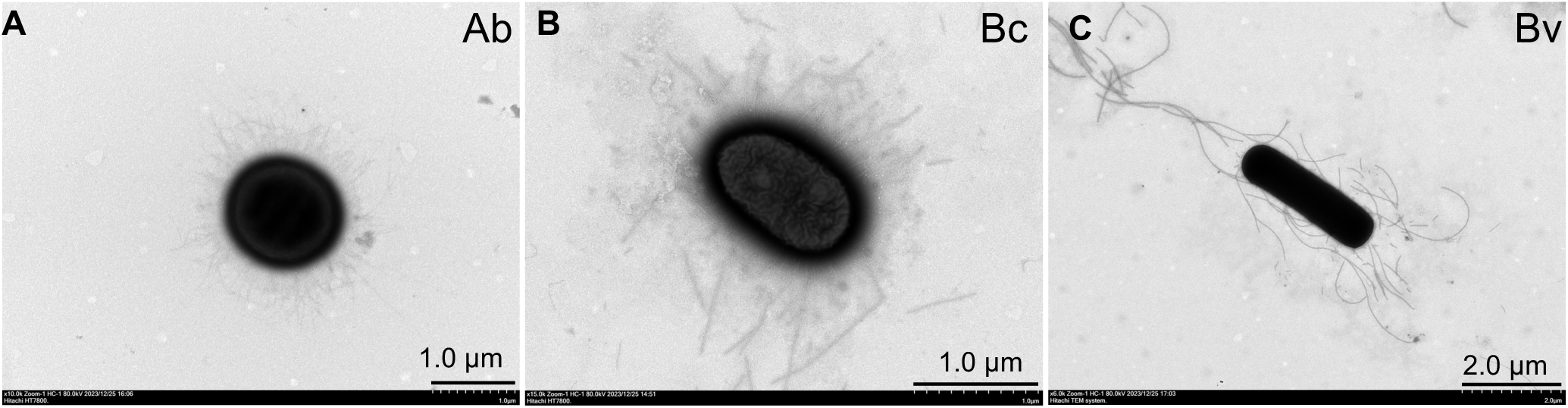
The TEM figures of Ab (A), Bc (B) and Bv (C). Scale bar showed in the figures.

**FIG S2.**
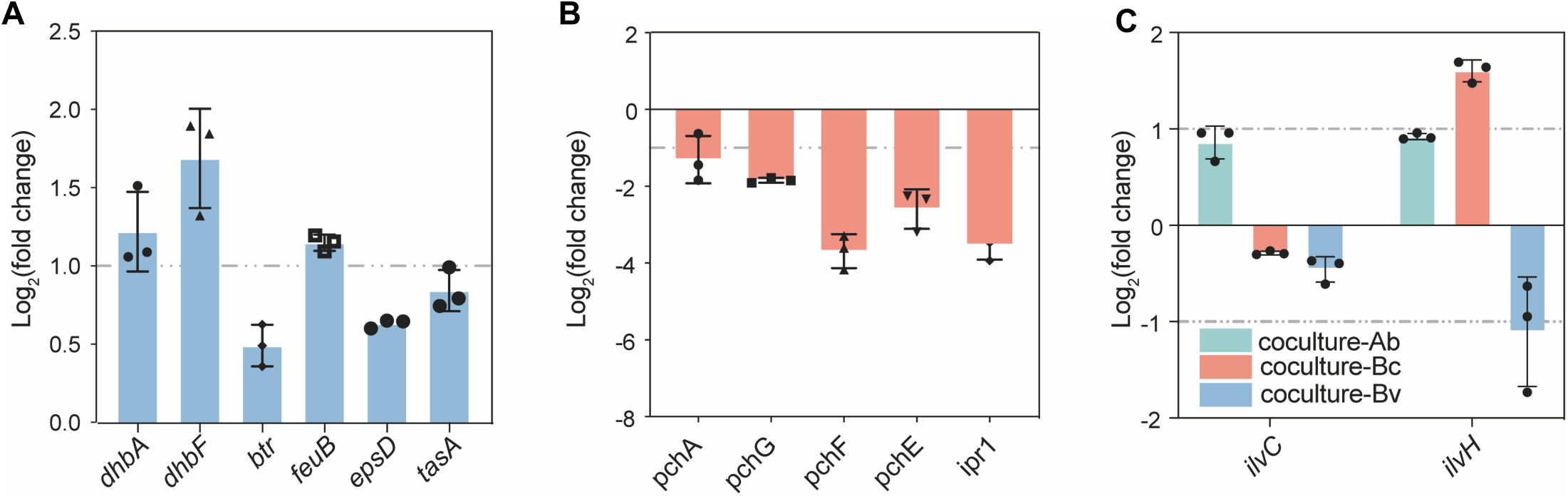
Transcriptome validation by qPCR. **(A)** Relative expression of genes involved in bacillibactin biosynthesis (*dhbA, dhbF and btr*), iron uptake (*feuB*) and biofilm matrix production (*epsD, tasA*) of Bv in tri-species biofilm compared with monoculture. **(B)** Relative expression of genes involved in siderophore biosynthesis in Bc in tri-species biofilm compared with monoculture. **(C)** Relative expression of *ilvC* and *ilvH* genes of the three isolates in tri-species biofilm compared with monoculture. Data presented are the mean ± s.d. (n = 3).

**FIG S3.**
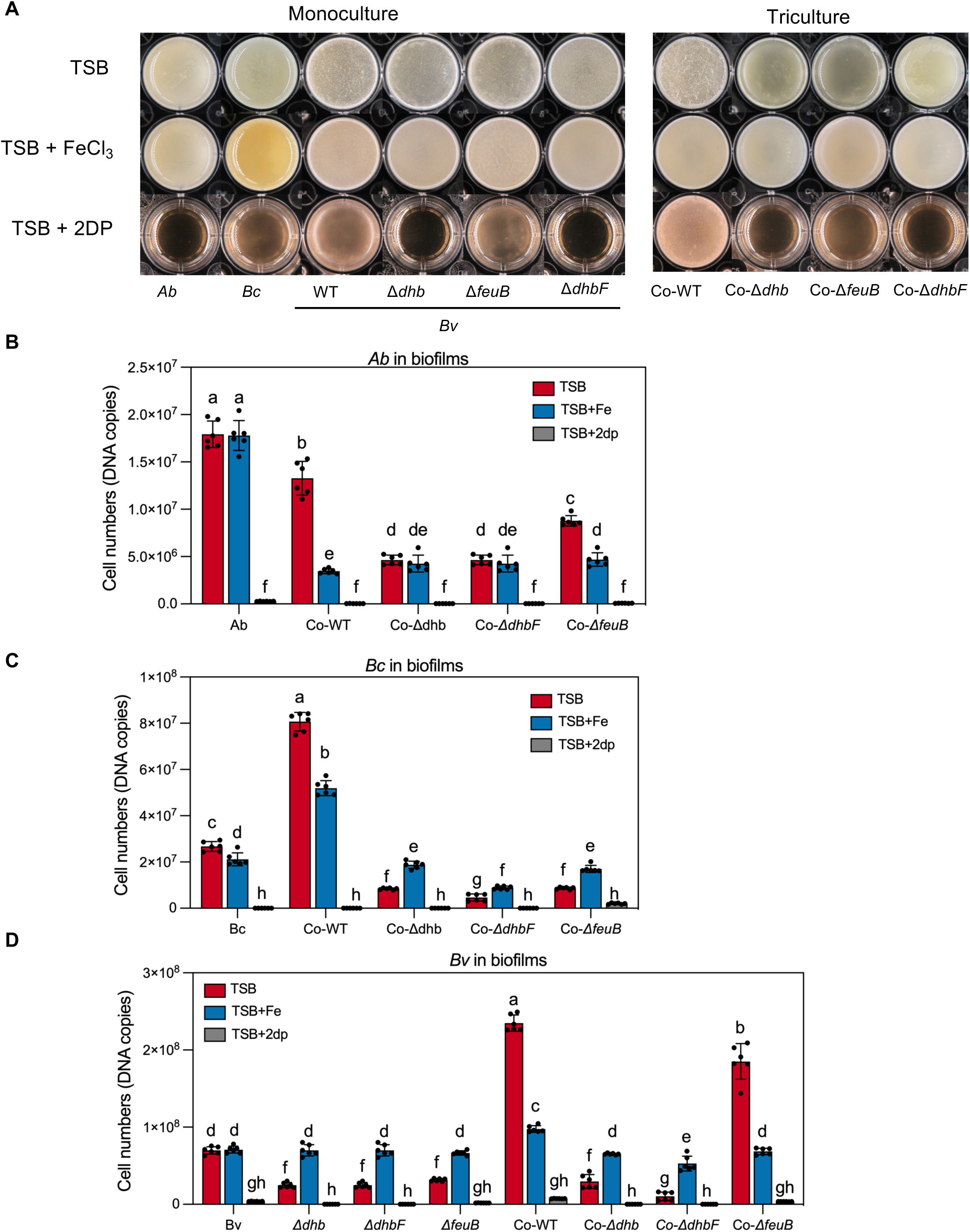
Biofilm phenotype and cell numbers quantification. **(A)** The biofilms formation in different treatments. **(B-D)** Cell numbers of *Ab*, *Bc and Bv*. Data presented are the mean ± s.d. (n = 6). Different letters indicate statistically significant (*p* < 0.05) differences according to one-way ANOVA followed by Tukey’s HSD post-hoc test.

**FIG S4.**
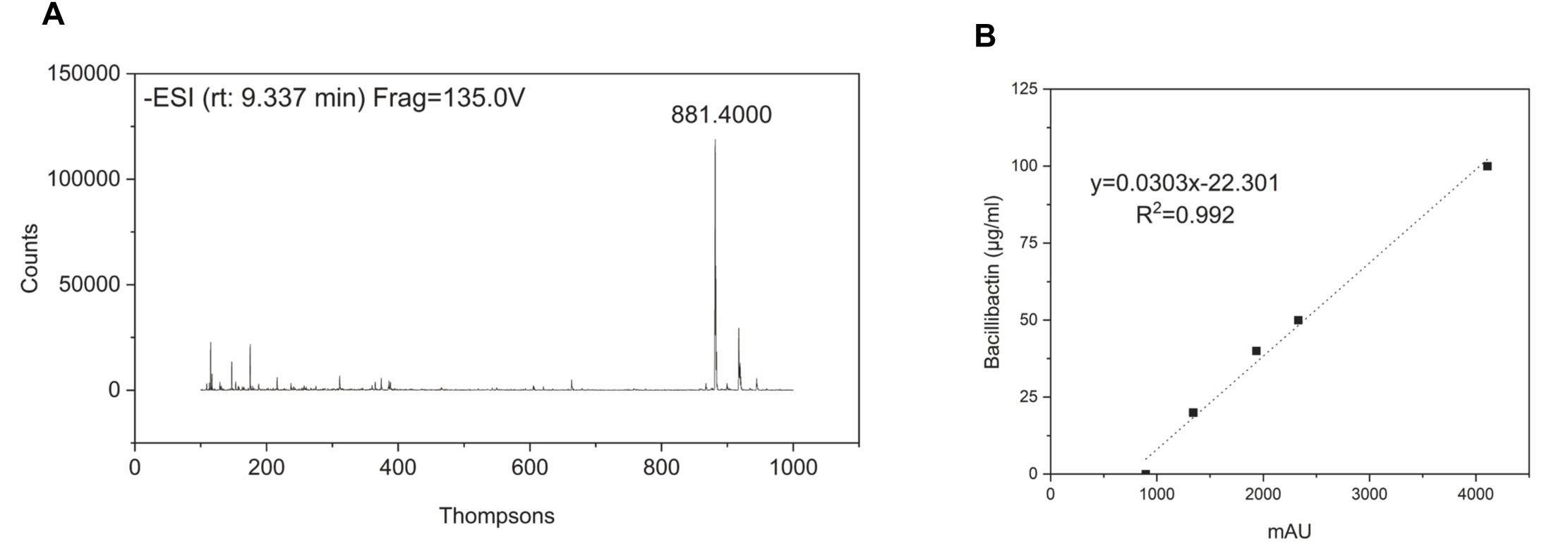
Analysis of bacillibactin. **(A)** LC-MS data of bacillibactin. **(B)** The standard curve of bacillibactin by HPLC.

**FIG S5.**
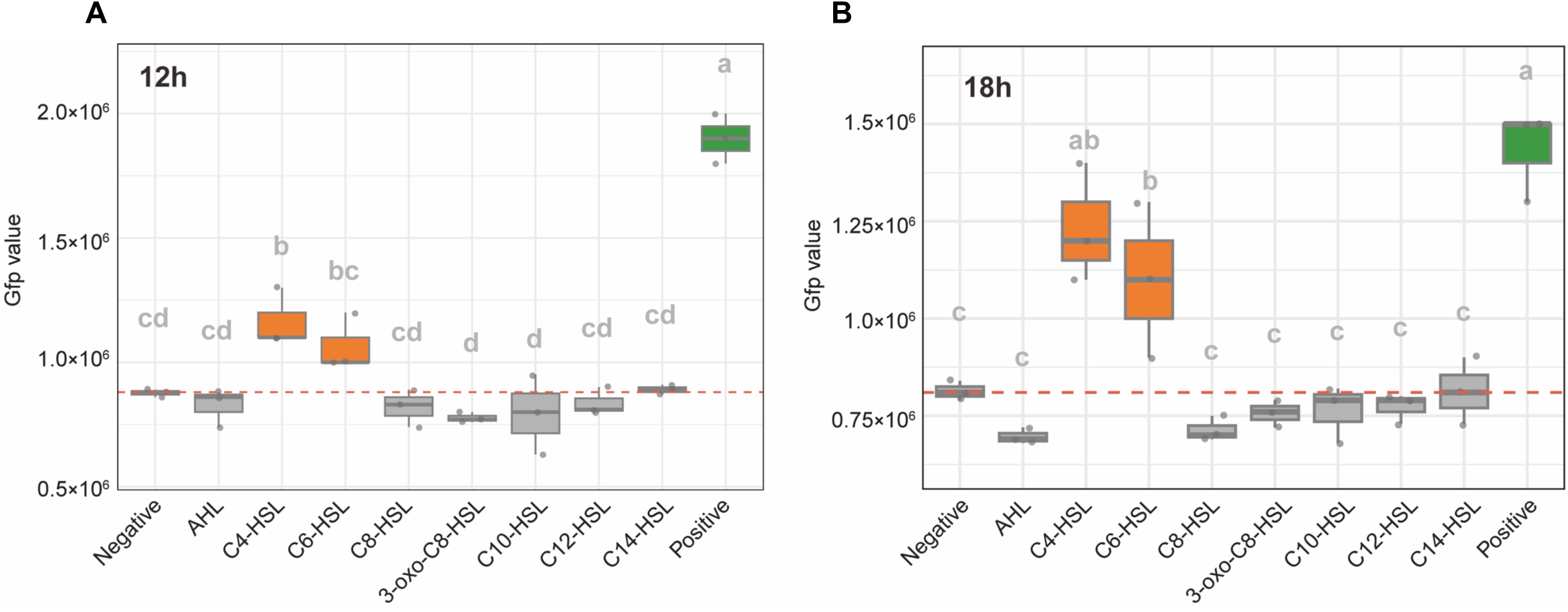
The *dhb* gene expression of Bv treated with different chemicals at 12h (A) and 18h (B). Negative: Bv *pdhb-gfp* alone. Positive: 50% of *Bc* supernatant with medium. Data presented are the mean ± s.d. (n = 3). Different letters indicate statistically significant (*p* <0.05) differences according to one-way ANOVA followed by Tukey’s HSD post-hoc test. Standard chemical list: Acetyl-L-Homoserine lactone (AHL), N-Butanoyl-L-homoserine lactone (C4-HSL), N-Hexanoyl-L-homoserine lactone (C6-HSL), N-Octanoyl-L-homoserine lactone (C8-HSL), N-(3-Oxooctanoyl)-DL-homoserine lactone (3-oxo-C8-HSL), N-decanoyl-L-Homoserine lactone (C10-HSL), N-Dodecanoyl-L-Homoserine Lactone (C12-HSL), N-Tetradecanoyl-DL-Homoserine Lactone (C14-HSL).

**FIG S6.**
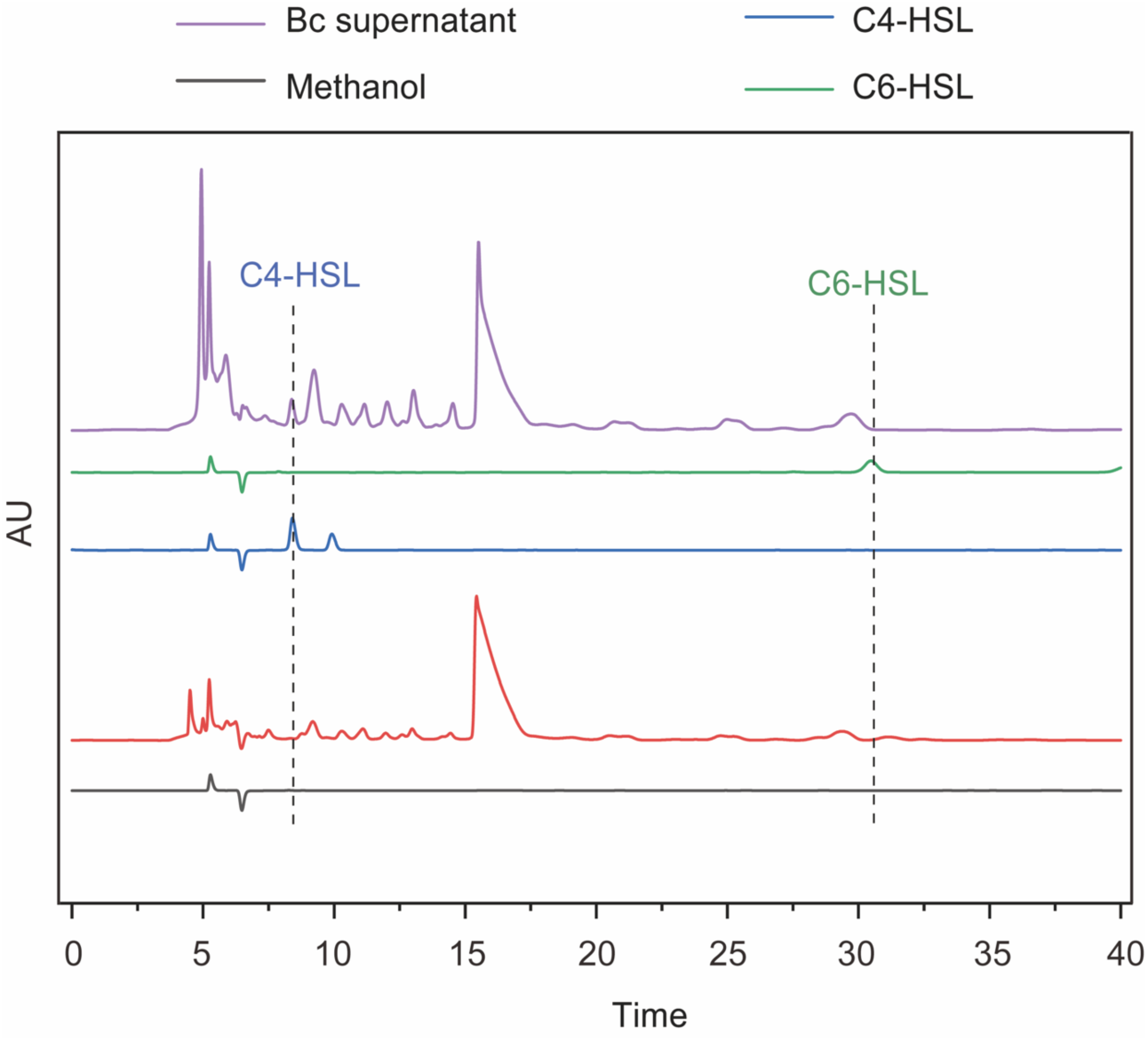
HPLC analysis Bc supernatant compared with C4-HSL and C6-HSL. The peak time of C4-HSL (N-Butanoyl-L-homoserine lactone) is 8.35 min The peak time of C6-HSL (N-Hexanoyl-L-homoserine lactone) is 30.50 min.

**FIG S7.**
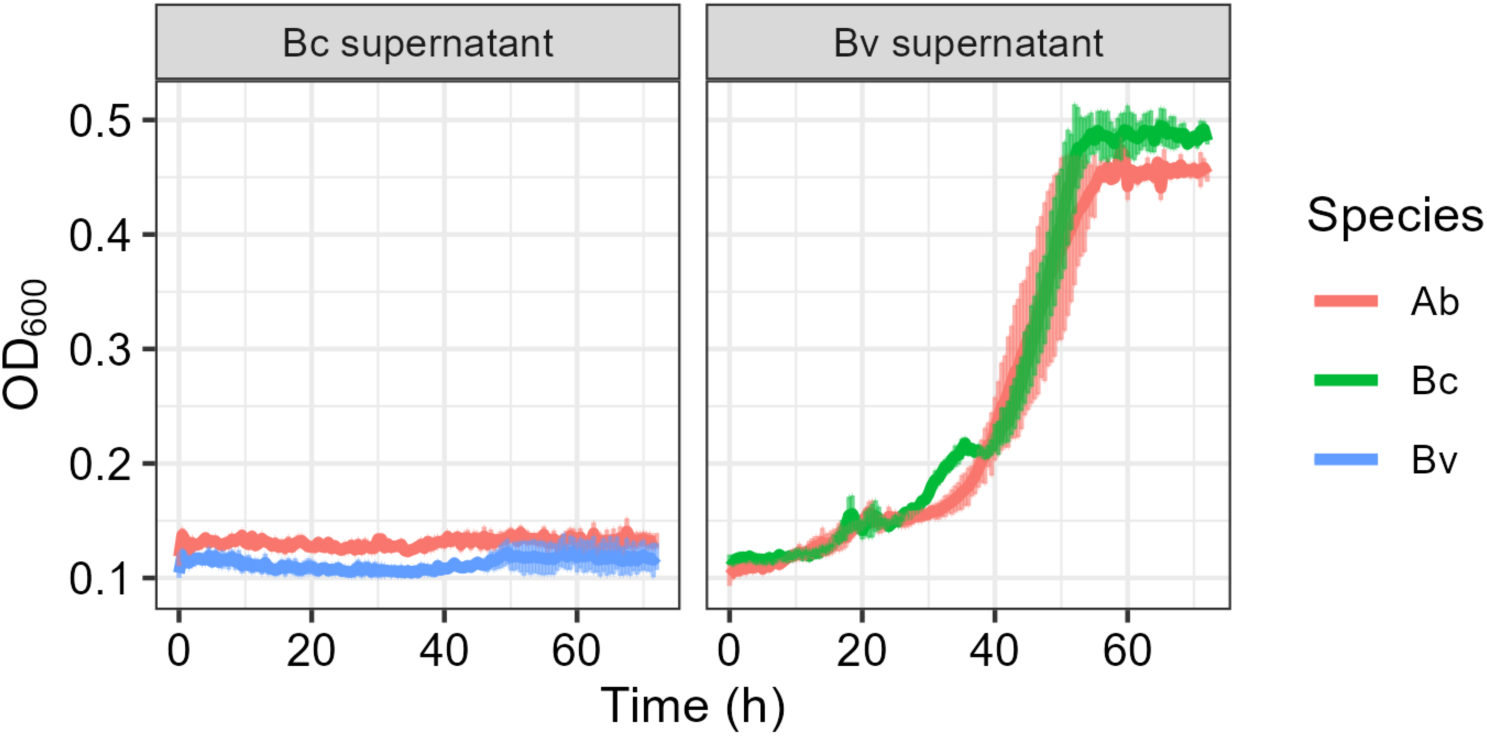
Growth of *Ab*, *Bc* and *Bv* in spent medium of either *Bc* or *Bv.* Data presented are the mean ± s.d. (n = 5).

**FIG S8.**
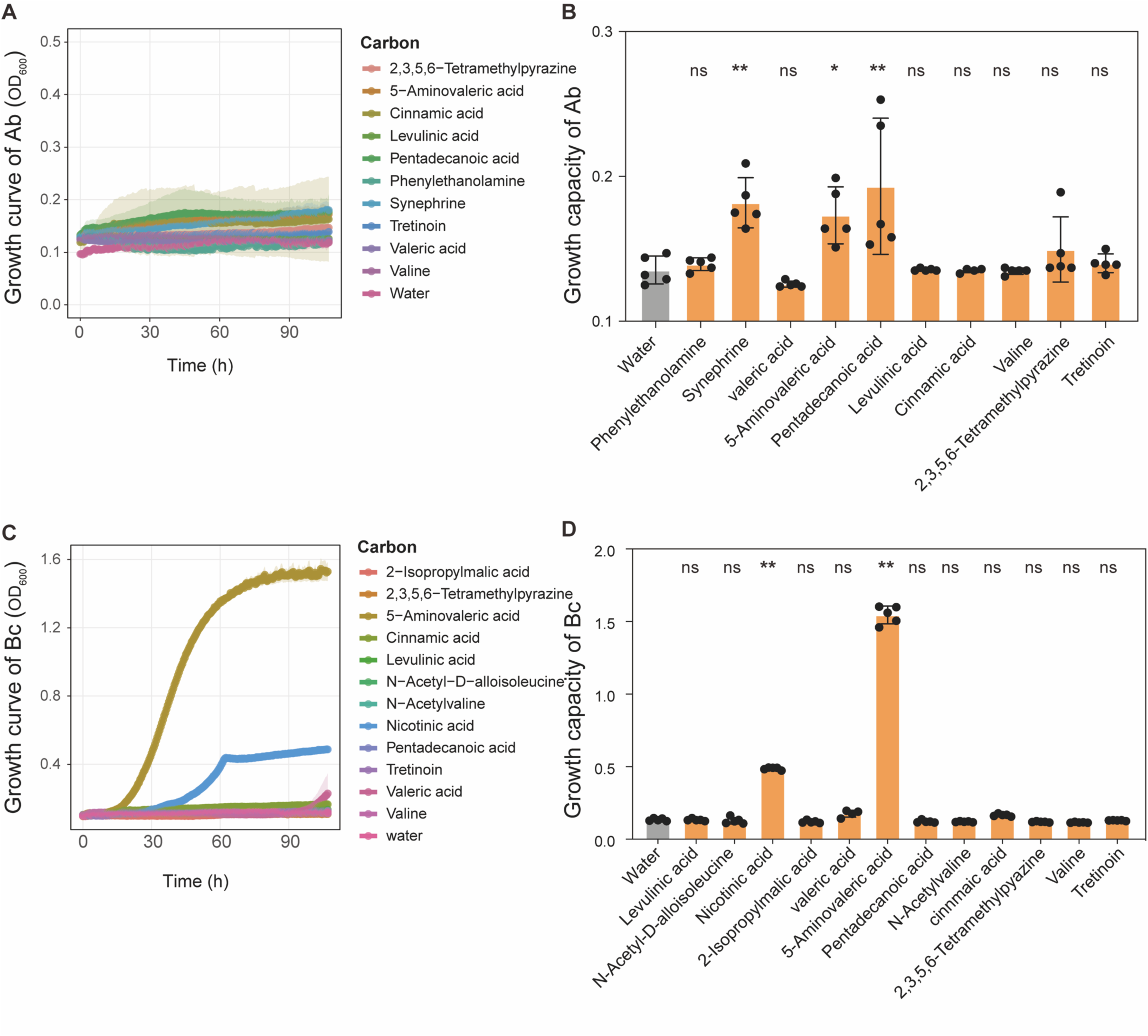
Metabolome results validation by growth curve. **(A&B)** Growth curves of Ab and growth capacity in M9 medium with corresponding compounds as sole carbon sources. **(C&D)** Growth curves of *Bc* and growth capacity in M9 medium with corresponding compounds as sole carbon sources. Growth capacity is the maximum population size. The value is the maximum OD_600_. Data presented are the mean ± s.d. (n = 5). Different letters indicate statistically significant (*p* < 0.05) differences according to one-way ANOVA followed by Tukey’s HSD post-hoc test.

